# Evolution of Mendelian dominance in gene regulatory networks associated with phenotypic robustness

**DOI:** 10.1101/2021.01.11.426187

**Authors:** Kenji Okubo, Kunihiko Kaneko

## Abstract

Mendelian inheritance is a fundamental law of genetics. Considering two alleles in a diploid, a phenotype of a heterotype is dominated by a particular homotype according to the law of dominance. This picture is usually based on simple genotype-phenotype mapping in which one gene regulates one phenotype. However, in reality, some interactions between genes can result in deviation from Mendelian dominance.

Here, by using the numerical evolution of diploid gene regulatory networks (GRNs), we discuss whether Mendelian dominance evolves beyond the classical case of one-to-one genotype-phenotype mapping. We examine whether complex genotype-phenotype mapping can achieve Mendelian dominance through the evolution of the GRN with interacting genes. Specifically, we extend the GRN model to a diploid case, in which two GRN matrices are added to give gene expression dynamics, and simulate evolution with meiosis and recombination. Our results reveal that Mendelian dominance evolves even under complex genotype-phenotype mapping. This dominance is achieved via a group of genotypes that differ from each other but have a common phenotype given by the expression of target genes. Calculating the degree of dominance shows that it increases through the evolution, correlating closely with the decrease in phenotypic fluctuations and the increase in robustness to initial noise. This evolution of Mendelian dominance is associated with phenotypic robustness against meiosis-induced genome mixing, whereas sexual recombination arising from the mixing of chromosomes from the parents further enhances dominance and robustness. Owing to this dominance, the robustness to genetic differences increases, while the optimal fitness is sustained up to a large difference between the two genomes. In summary, Mendelian dominance is achieved by groups of genotypes that are associated with the increase in phenotypic robustness to noise.

**Author summary:** Mendelian dominance is one of the most fundamental laws in genetics. When two conflicting characters occur in a single diploid, the dominant character is always chosen. Assuming that one gene makes one character, this law is simple to grasp. However, in reality, phenotypes are generated via interactions between several genes, which may alter Mendel’s dominance law. The evolution of robustness to noise and mutations has been investigated extensively using complex expression dynamics with gene regulatory networks. Here, we applied gene-expression dynamics with complex interactions to the case of a diploid and simulated the evolution of the gene regulatory network to generate the optimal phenotype given by a certain gene expression pattern. Interestingly, after evolution, Mendelian dominance is achieved via a group of genes. This group-based Mendelian dominance is shaped by phenotype insensitivity to genome mixing by meiosis and evolves concurrently with the robustness to noise. By focusing on the influence of phenotypic robustness, which has received considerable attention recently, our result provides a novel perspective as to why Mendel’s law of dominance is commonly observed.

## Introduction

Mendelian inheritance is a keystone of genetics. Mendel’s law [1] concerns binary traits and consists of three laws: the law of gene segregation, the law of independent assortment, and the law of dominance. From a modern viewpoint, the law of segregation is explained by meiosis and the law of independent assortment by the “independent” expression of different genes [2].

The law of dominance assumes a pair of alleles, which involves a nonlinear interaction leading to an epistasis between the genes. A binary trait (phenotype) is given by A and a. If the two genes in the given alleles are homo and *AA* (*aa*), the trait is A (a). For a heterozygote, i.e., *Aa*, the trait is A if A is dominant. Indeed, by creating a pure line that has *AA* and *aa*, Mendel showed that F1, a hybrid of *AA* or *aa*, expresses only the dominant character A, whereas the next generation from F1 shows character A or a according to a 3:1 ratio.

In reality, there are many genes in different loci. Nevertheless, their independence is assumed in the standard Mendelian inheritance. Even if there are many genes, when expression is independent, each trait is determined by the alleles according to simple genotype-phenotype mapping [3]. However, in general, multiple genes in different loci often interact with each other, which can result in genotype-phenotype mapping. Recent studies have demonstrated that genes often constitute a gene regulatory network (GRN). Although it is possible to establish a pure line for the alleles that corresponds to a specific trait, there are other ‘hidden genes’ that may influence the trait. Consequently, the law of dominance can be violated, with the traits of F2 deviating from the Mendelian ratio of 3:1, as is measured by the interaction deviation [4, 5].

Furthermore, the phenotype, even in the homozygotous case, can be perturbed by noise [6, 7]. This can also result in deviation from the Mendelian ratio.

In contrast, the possibility of generalizing Mendelian dominance beyond the simple single-gene case has been discussed previously. Fisher posited that Mendelian dominance could arise because of evolutional benefits concerning robustness [8]. Alternatively, Wright argued that metabolic stability could lead to dominance [9]. However, to date, both arguments remain hypothetical, and any quantitative relationship linking the robustness in gene expression or metabolic dynamics to Mendelian dominance remains elusive.

Then, the degree of deviation from Mendel’s law of dominance by the interaction among genes and stochasticity in the gene expression need to be studied, in relationship with the robustness. In general, such degree as well as the robustness depends on the nature of gene expression dynamics governed by the GRN, which is shaped through the course of evolution. Indeed, in the haploid case, extensive studies have been conducted on the nature of gene expression dynamics governed by gene-gene interactions under the influence of stochasticity. The evolution of genotype-phenotype mapping has been investigated numerically using the GRN model, thus elucidating the evolution of robustness to noise and mutations [10–13]. Therefore, it is important to extend these studies of gene expression dynamics to the diploid case [14, 15].

Here, by simulating the evolution of diploid GRNs, we seek to determine whether the degree of Mendelian dominance evolves beyond the classical case of one-to-one genotype-phenotype mapping. Specifically, we examine whether complex genotype-phenotype mapping can achieve Mendelian dominance through the evolution of the GRN with interacting genes. We adopt a computational model of the GRN and adapt it to the diploid case, introducing a genetic protocol to account for sexual recombination and meiosis in the diploid. Note that, in the diploid case, proteins are transcribed from the two alleles. As such, the expression dynamics are a result of the superposition of two GRNs through meiosis segregation, sexual recombination, and mutation. Next, the results of the evolution simulation demonstrates the evolution of Mendelian dominance via a group of interacting genes in the GRN. The condition for the evolution of Mendelian dominance and its possible relationship with robustness to noise and mutation is explored. Additionally, the importance of meiosis-based genome mixing in establishing this dominance is revealed. Finally, we discuss the possible connection of this group-based Mendelian dominance to experimental observations.

## Results

### Evolution simulation based on a diploid GRN model

First, we introduce the theoretical model of the GRN. Here, we consider the gene expression level *x_i_* (−1 < *x_i_* < 1) for the genes *i* = 1, …, *k*. Each gene *i*(= 1, 2, …, *N*) has an expression level *x_i_*(*t*) at time step *t*. In the model, *x_i_* is scaled in order that it takes a value between −1 and +1 (instead of 0 and 1) to emphasize that the two states, i.e., expressed (+1) or non-expressed (−1), are symmetric. Each gene interacts with others and itself, with the interaction of the *j*-th gene with the *i*-th gene is described by the matrix *J_ij_* [3, 11, 12, 16, 17]. *J_ij_* can take three values: +1, −1, and 0, which represent the activation of gene *i* by gene *j*, the corresponding inhibition, and no interaction, respectively.

We adopt a discrete-time model [3, 6, 18, 19], in which the expression level *x_i_*(*t* + 1) in the next time step is determined by 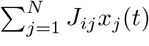 represents the GRN. A small Gaussian noise, *N* (0*, σ*), is added to the dynamics to account for stochasticity in the gene expression. By using the sigmoid function *f* (*x*) = tanh(*βx*) with large *β*(= 100), the expression dynamics are given by

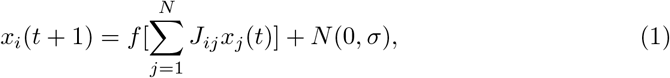

which is the discrete-time version of the continuous-time model for gene expression dynamics [10, 17]. The initial expression levels {*x_i_*(0)} take a value of +1 or −1 for each gene and are fixed for all individuals over generations. After a certain time, *T*, *x_i_*(*t*) reaches a fixed point for most cases, where *x_i_* is ±1 depending on the structure of the network, *J_ij_*. Notably, after the evolution of *J_ij_*, *x_i_*(*t*) always reaches a fixed point. Thus, after sufficient time steps (*T*), the expression pattern *x_i_*(*t*) determines the phenotype. Note that there are some cases where the expression pattern *x_i_*(*t*) cannot reach a stable state; however, these are rare.

Diploids have two genomes that correspond to two matrices, 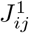 and 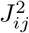 [14, 15, 19, 20]. The gene expression dynamics of diploids are determined by the sum of these matrices such that the dynamics are given by modifying *J_ij_* to 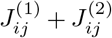 in Eq. 2:

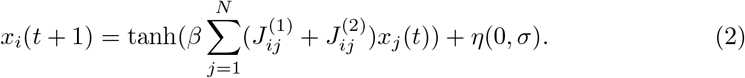

Genetic changes take three main forms: asexual, meiosis only (i.e., without recombination), and the standard sexual protocol, which includes both meiosis and recombination. In this paper, we term these, asexual, meiosis only (MO), and meiosis and recombination (MR). In the asexual protocol, the two genomes are not mixed, i.e., 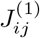 and 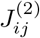 are copied. The other two protocols involve sexual reproduction. In MO, one genome from parent−1 becomes a new 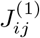 (gamete) and one genome from parent−2 becomes a new 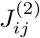. In MR, two genomes from parent−1 (parent−2) are mixed by recombination, to provide a gamete, that is, a new 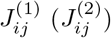. In recombination, 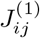 and 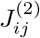 are mixed by a row vector in both parents to produce a gamete. Recall that mutation is included as a change in the matrix element *J_ij_* with 0 or ±1 in all three protocols. A fourth protocol, involving recombination only(RO), is included as a reference to elucidate whether meiosis or recombination is more important for Mendelian dominance (see Methods for details).

To obtain fitted phenotypes, 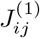 and 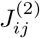 were selected as parent(s). Here, fitness is defined by how closely the expression pattern, {*x_i_*}, matches a prescribed target. (See Methods for the details of target and selection procedure.)

### Evolution of fitness

The simulated evolution of the fitness is shown in Fig. 1, with the evolution of the average fitness of the population plotted for the asexual, MO, and MR protocols. In each case, fitness increased beyond 90% of the maximum, within 500–1000 generations (Fig. 1), with MR and MO exhibiting the fastest and slowest increase, respectively.

**Fig 1.**
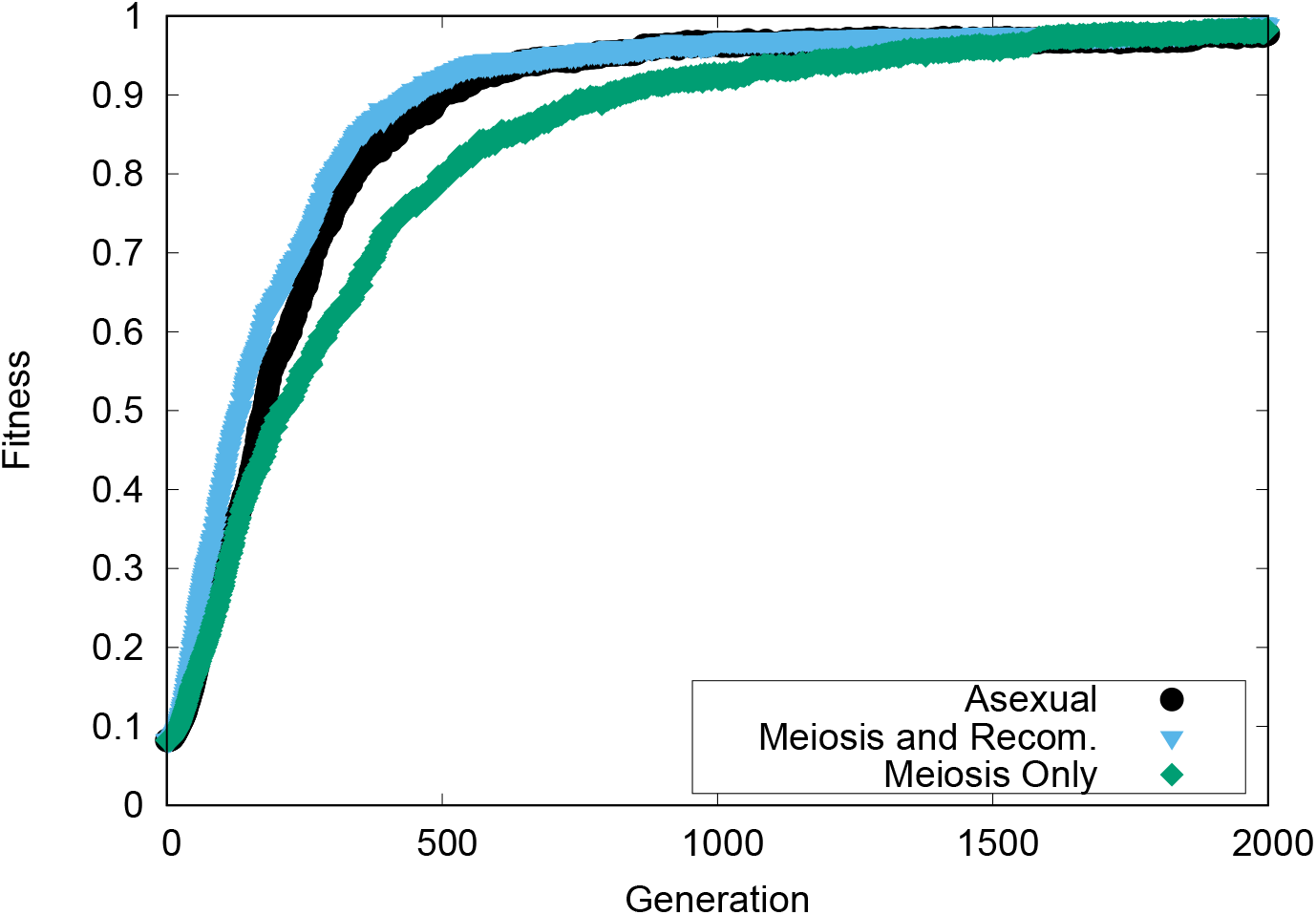
**Fitness as a function of evolutional generation** for the asexual (black line), MR (blue line), and MO (green line) protocols. The mutation rate per edge = 4 × 10^*−*5^ and the strength of noise = 0.01. The increase in fitness was saturated within 2000 generations.

Figure 2 shows the mutation rate dependence of the fitness. For each protocol, as the mutation rate increases, the increase in fitness stops. A drop begins at a mutation rate of approximately 10^*−*3^. For MR, this decline is slightly suppressed and is initiated at a slightly higher mutation rate. This decline is caused by the loss of information at higher fitness values owing to the accumulation of mutations in the network, as discussed in the context of error catastrophe [21]. Although, the recombination process in MR slightly suppresses this error catastrophe, the effect is not significant.

**Fig 2.**
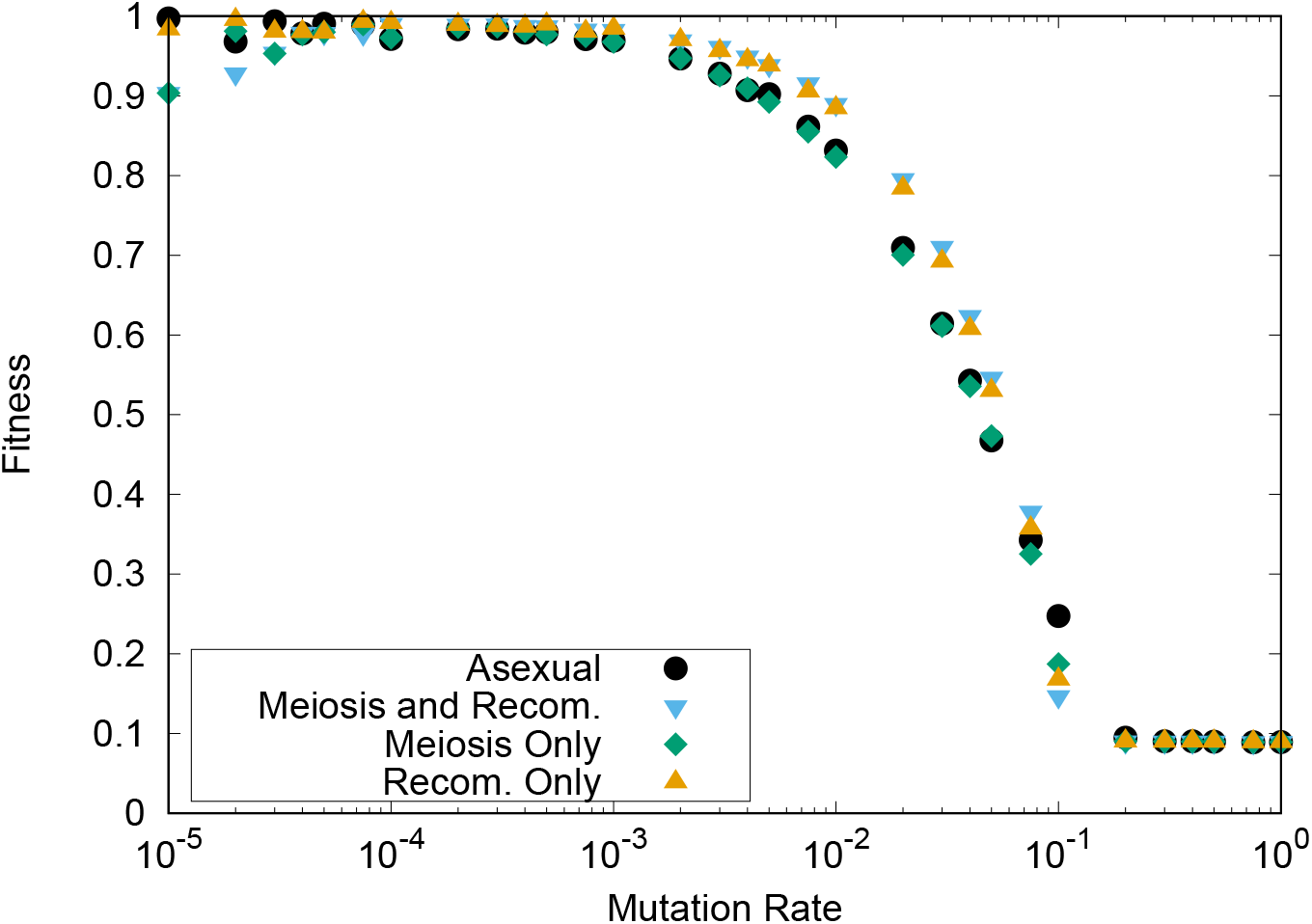
Mutation rate dependence of the average fitness. The average fitness was computed for 100 individuals representing the last generation in the evolution of over 10000 generations for the asexual (black circles), MR (blue triangles), and MO (green diamonds) protocols. The noise amplitude *σ* was 0.0001. For mutation rates exceeding 10^*−*1^, error catastrophe occurs, which prevents the average fitness from rising through evolution over 10000 generations. The recombination only reference case is also plotted (orange triangles).

### Group-based Mendelian dominance

Next, we investigated if and how Mendelian dominance evolves. As mentioned in the Introduction, the inter-gene interactions result in deviation from the 3:1 ratio of Mendelian dominance [4, 5]. Here, we rename the phenotypes, +1 and −1 with + and − with normal font. Corresponding genotypes are written in italic like + and −.

We create a pure line for a homozygous case in which two genomes have the same *J_ij_* in the diploid [22]. There are a variety of matrices (genes) that produce + (−) for a specific *x_i_*, which are given by *+1+1, +2+2, +3+3*, …, *+n^+^+n^+^* (*−1−1, −2−2, −3−3*, …, *−n^−^−n^−^*). From the + (*+1, +2*, …, *+n^+^*) and - (*−1, −2*, …, *−n^−^*) sets, we generate *N*_sample_ homozygotes, *++* or *− −*, and 2*N*_sample_ heterozygotes, *+-*, whose phenotypes can be examined. Then, we count the number of each case, denoted by 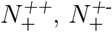, and 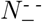, respectively (Here, we assumed that the number of 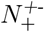 is larger than 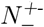 and that + is dominant. We rename + and − if 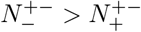) when Mendel’s law of dominance is perfect, 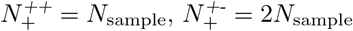, and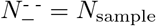. However, in general, the genes are not independent, with the groups + and − chosen as a set of genotypes that produce each trait such that each fraction deviates from a ratio of 1:2:1. Therefore, we define the group Mendelian ratio (GMR) as the fraction 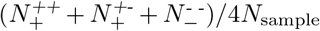. The deviation of the GMR from unity corresponds to the interaction deviation in quantitative genetics [4, 5] (see the Methods for the detailed algorithm).

### Condition for the increase of the group Mendelian ratio

First, we studied the dependence of the GMR on the mutation rate (Fig. 3). In the MO and MR protocols, the average value of the GMR almost exceeds 0.8 for mutation rates less than 10^−3^ (note that this is an average for all genes *i*: For *x_i_* of some genes GMR reaching 1). For mutation rates exceeding 10^−3^, the average GMR decreased, and the value for MR was higher than that for MO. In contrast, for the asexual case, the GMR remained at a much lower level (∼ 0.65) and peaked at approximately 0.75. Note that, in the case of recombination without meiosis, GMR remained at the same low level as in the asexual case.

**Fig 3.**
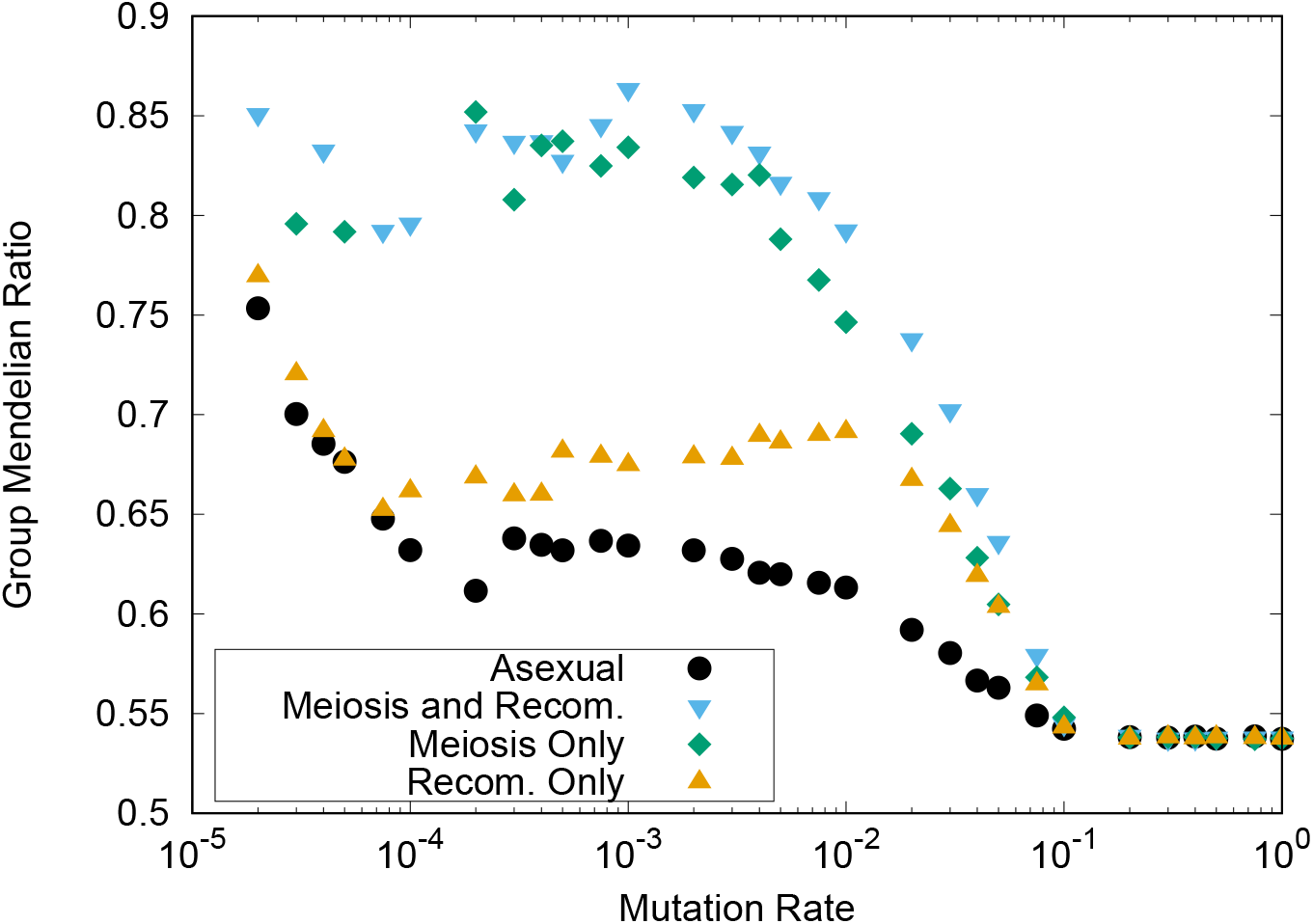
Group Mendelian ratio as a function of mutation rate. The GMR at the 10^4^-th generation is shown for the asexual (black dots), MR, (blue triangles), and MO (green diamonds) protocols. The noise amplitude was 0.0001. Data for the recombination only reference case is also plotted (orange triangles).

Recall that GMR is 0.5 if all the GRNs are random. If the GMR is sufficiently high, group-level Mendelian dominance occurs, which we term “group Mendelian dominance” (GMD). Specifically, we regard GMD to be achieved when the average GMR in the population *>* 0.8. Therefore, GMD is realized for the MO and MR protocols, i.e., when meiosis is considered.

### Correlation between group Mendelian ratio and robustness to noise

#### Phenotypic fluctuation and the group Mendelian ratio

We have shown that the mutation-rate dependence of the GMR for the MR and MO cases is highly correlated with that of the fitness, as evident upon comparing Fig. 2 with Fig. 3. Therefore, we plotted the correlation between the GMR and the fitness in Fig. 4a. As shown, the two exhibit strong correlation. Nevertheless, the plot in Fig. 4a Fig. 4c shows three branches depending on the three different noise levels adopted here.

**Fig 4.**
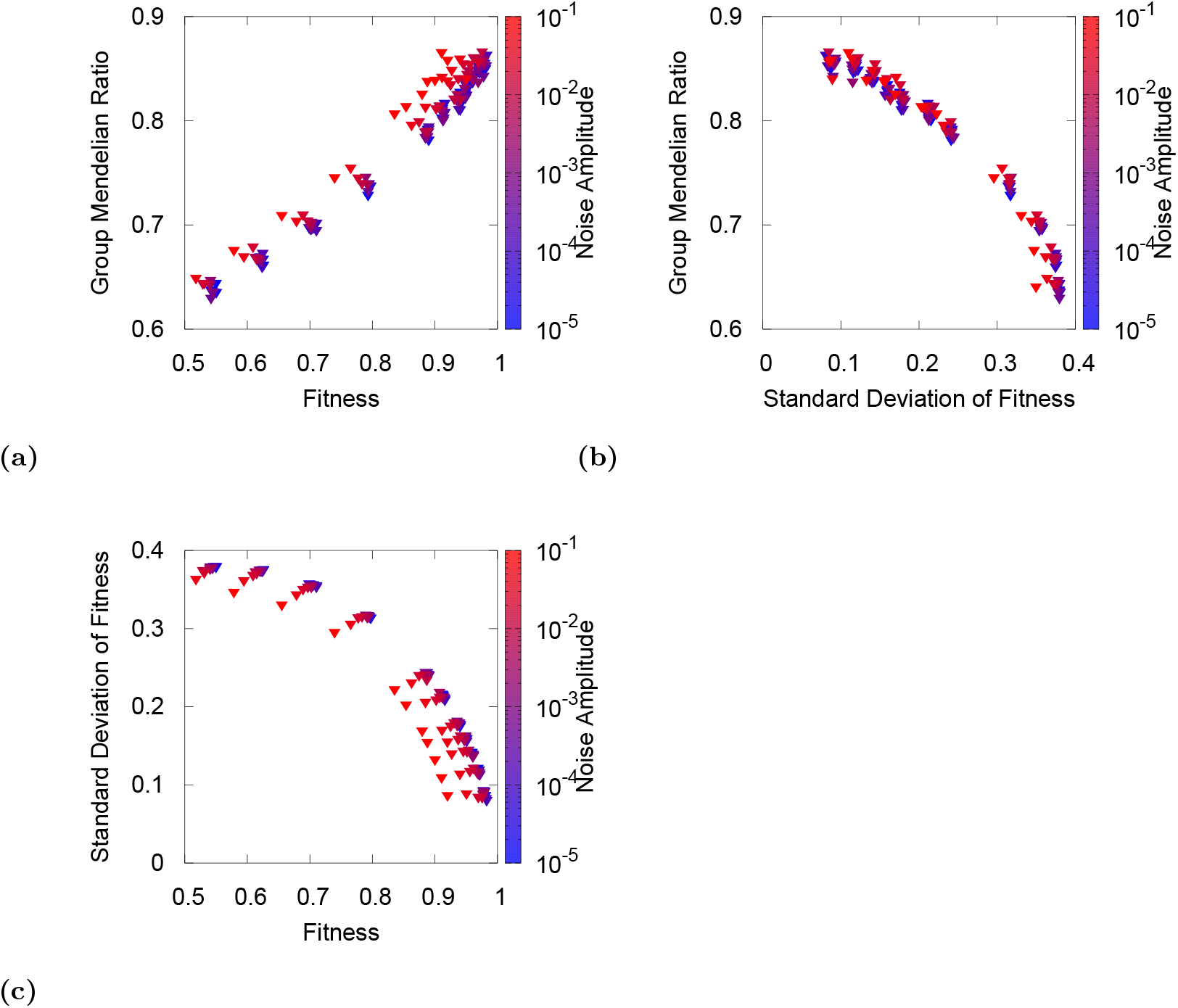
Correlation between the GMR, the average fitness, and the standard deviation of the fitness. Each variable was computed as the average over populations of 10 generations before 10^4^ generations, with the evolution having reached a steady state. The GMR was computed as the value of the 10^4^-th generation for the MR case. The points were obtained across different mutation rates from 10^−5^ to 10^−1^, as indicated by the color of the data, while the noise level satisfied 0.001 ≤ *σ* < 0.075. (a) GMR as a function of the average fitness, (b) GMR as a function of the standard deviation of the fitness, (c) standard deviation of the fitness as a function of the average fitness. Note that (a) and (c) each have three branches corresponding to different noise levels, whereas (b) does not, implying that the GMR is dominantly correlated with the fluctuation of the fitness, rather than its average.

As the noise level varies, the variances of the phenotype and fitness also vary. Therefore, to examine the relationship between the GMR and noise-induced phenotypic variances, we plotted the GMR against the standard deviation of the fitness. In this case, the correlation is more prominent, with the data not exhibiting noise level-related branches. All data from different mutation rates and noise levels were well fitted by a single curve, as shown in Fig. 4b. Indeed, the difference between Figs. 4a and 4b also is further explained by the plot of the fitness variance against the average fitness (Fig. 4c), which, again, shows three branches. Together, these three plots indicate that the GMR shows a better correlation with the fitness variance than the fitness itself.

### Robustness to initial noise and the GMR

In general, the smaller the fluctuations, the less sensitive the corresponding state is to perturbations. As a consequence, the phenotype, specifically the fitness, becomes more robust [10, 17]. Therefore, we investigated the correlation between the GMR and the robustness of the fitness to noise (Fig. 5). We considered the robustness to noise with respect to the initial expression levels {*x_i_*(0)}, as they influence the later gene expression dynamics and, thus, the robustness to initial condition correlates strongly with that to later purturbations. Here, we define the noise robustness as the ratio of the fitness under a given initial noise to the original noiseless fitness value.

**Fig 5.**
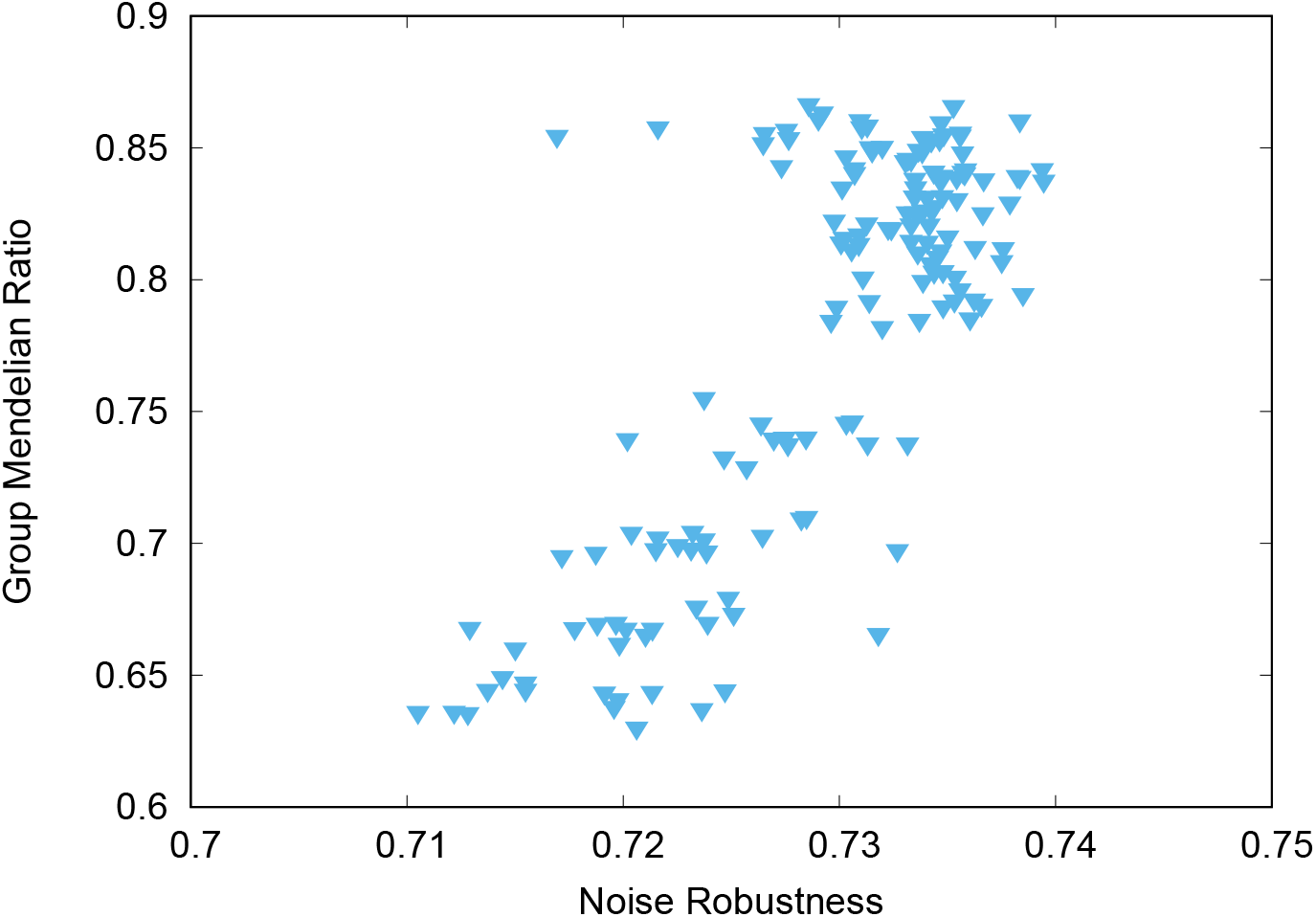
Group Mendelian ratio as a function of the robustness to noise. The GMR was computed following the same procedure as in Fig. 4. To calculate the noise robustness, perturbed initial conditions were chosen in which the expression of each gene was switched, from which the fitness and, subsequently, the average fitness were computed. The noise robustness was defined as the ratio of the average fitness for the perturbed initial condition to the original. We plotted for the MR case. The data correspond to the equivalent mutation rates and noise levels as used in Fig. 4.

As shown in Fig. 5, the GMR is correlated with the robustness to initial noise. As the robustness increases with the decrease in fluctuations, the negative correlation observed between the fitness fluctuations and the GMR in Fig. 4b can be attributed to the correlation between the robustness to noise and the GMR.

Note that an equivalent correlation between the GMR and the robustness to mutation or recombination is not observed. This is because robustness to mutation or recombination has already been achieved for diploids after the evolution progresses, and the robustness is no longer reduced.

### Group Mendelian ratio and distance between diploids

As the phenotype is shaped according to genotype-phenotype mapping by the GRN, it is also important to study the correlation of the GRN with genetic variance. In diploids, the distance between two genomes can provide a measure of the genetic variation. Fig. 6 shows the relationship between the GMR and the genetic distance between two genomes in a diploid. As shown, the GMR begins declining at a certain distance, with the fitness also dropping. Additionally, if the genetic difference between the two genomes is too high, the GMD is not sustained.

**Fig 6.**
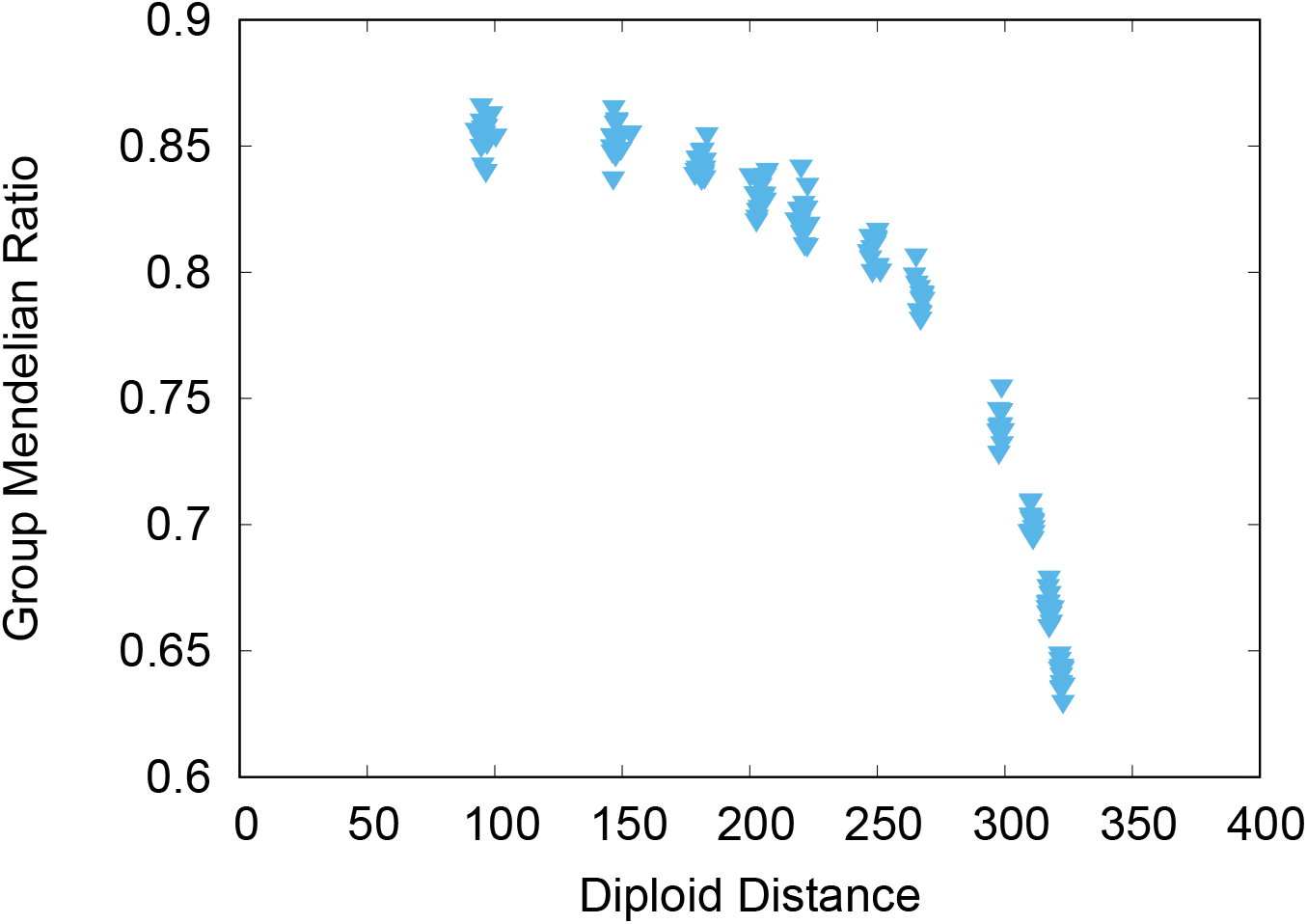
Group Mendelian ratio as a function of the distance between the two genomes. The distance was computed as the Euclidean distance between the matrix elements in two genomes, 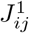 and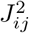. All data were obtained from the evolved GRNs for the MR case shown in Fig. 4. We simulated GRNs with 20 genes, such that the number of matrix elements, that is, the maximal Hamming distance (i.e., the number of different elements) is 20 × 20 = 400. Consequently, the GMR (and the fitness) maintains a high value until half the elements have been changed.

In Fig. 6, the GMR can exceed 0.8 when the distance is under 250. Note that the total number of matrix elements (genes) is 20×20, which ensures that the fitness or GMR is maintained, even if over half the elements are different between two genomes. Furthermore, high robustness is achieved to maintain fitness against meiosis. This implies that the fitness of the diploid (with meiosis) is remarkably robust against genetic differences between the two genomes. Recall that meiosis mixes genomes in the diploid. However, the fitness should be robust to such mixing, which explains the insensitivity of the fitness.

## Discussion

In this study, we have investigated a diploid GRN model by extending the model described by Omholt et al. [14]. In our model, two GRNs work concurrently to yield the phenotype as a set of gene expressions. By evolving the GRN with mutations, meiosis, and recombination, we have shown that Mendelian dominance evolves. This dominance is achieved via a group of assorted genotypes that share a common phenotype, which is determined by the expression of target genes. The degree of dominance increases (corresponding to the decrease of interaction deviation [4, 5]) through evolution, exhibiting a strong correlation with the decrease in phenotypic fluctuations and the increase in the robustness to initial noise. This evolution of Mendelian dominance is prominent in the presence of meiosis, which mixes two genomes, whereas under sexual recombination, the mixing of genomes from the parents further enhances this dominance and robustness. Mendelian dominance increases the robustness to genetic differences, and the fitness is maintained against large differences between two genomes.

For Mendelian dominance in individuals, the heterozygote advantage has been argued by Goldstein [23] and Poter et al. [24]. Moreover, the relationship between Mendelian dominance and mutational robustness has been investigated by Bagheri and Wagner [25], whereas the influence of ploidy or recombination on robustness has also been discussed [15, 19, 20].

Indeed, a possible connection between Mendelian dominance and robustness has been posited previously. Fisher discussed advantages of dominance for the evolutionary robustness of phenotypes [8], while Wright emphasized the relationship between dominance and stability in metabolic networks [9]. However, these discussions were primarily qualitative. Our study integrates these qualitative arguments to extract quantitative information by using gene expression dynamics and a quantitative definition of the GMR.

The degree of dominance introduced in our model follows the standard method of establishing pure lines and, thus, can be measured by proper experiments. For example, Hou et al. [26] tested the degree to which Mendel’s dominance law holds in yeast, while also measuring the phenotypic variation. Accordingly, the validity of the negative correlation between Mendelian dominance and the phenotypic fluctuation revealed in our theoretical models can be verified by measuring the degree of Mendelian dominance and the fluctuations across genes or between different strains.

In addition, Mendelian dominance in plants has been studied by Huber et al. [27] and Govindaraju [28]. Note that ploidy beyond diploidity is more common in plants. Nevertheless, extending our model to consider ploidy is straightforward, enabling the relationship between dominance and phenotypic fluctuations to be examined and testable predictions to be made.

For the simulation presented herein, we can compare the outcomes of different genetic protocols, namely, asexual, MO, and MR. Our analysis concludes that the existence of meiosis is essential to realize GMD. Although the fitness is only slightly changed from the MR case, the GMR (and diploid distance in Fig. S1) is reduced drastically relative to the MR and MO cases. Therefore, meiosis-induced genome mixing is essential for GMD. Genome mixing between diploids requires the GRN to ensure that high fitness is sustained following gene alterations in the other gene. This suggests that the phenotype between two alleles in the hetero case is identical to that in the “homo” case, leading to GMD. From a contemporary viewpoint [2], meiosis is often regarded as consistent with segregation in Mendel’s law. Based on our study, one can conclude that Mendelian dominance stems from Mendelian segregation.

## Materials and methods

### Fitness for evolution

After the expression pattern *x_i_*(*t*) reaches the stable state *x_i_*(*T*), the fitness is determined by the distance between the gene expression of the individual and the prescribed target pattern. This target pattern *p_i_* is defined for a part of the genes, namely the “target genes” *i* = 1, …, *M* (*< N*). Here, the phenotype is defined as the time average of *x_i_*(*t*) for the last 10 steps, 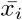. Accordingly, a stable expression pattern can be more advantageous than an oscillatory state. Setting the maximum fitness for, 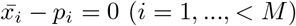 the fitness is defined as 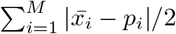. Furthermore, the selection pressure is set as *S*, such that the probability of obtaining the phenotype 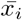 is proportional to 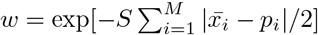.

### Genetic protocols

Using the fitness, a diploid GRN can be produced in the next generation. To introduce genetic variation, we adopted four genetic protocols: asexual, meiosis only (MO), meiosis and recombination (MR), and recombination only (RO).

In the asexual protocol, offspring are generated as a copy of a parent. One individual, *k*, is chosen as a parent with probability *w_k_/W*, where 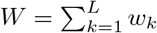. The matrices 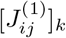 and 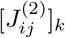 of this parent are copied for the next generation.

In MO, two individuals, *k*_1_ and *k*_2_, become parents with probabilities 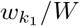 and 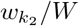, respectively; *k*_1_ and *k*_2_ must be different. Next, a gamete, 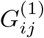, is produced from the diploid of one parent, *k*_1_. One of the two genomes in *k*_1_ becomes 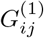, here. The other gamete, 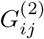, is produced from *k*_2_ following the same procedure. These two gametes, 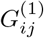 and 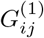 become 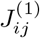 and 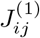 for an individual in the next generation.

In MR, genome mixing process is same as MO but gametes are generated by a different protocol including recombination. Forming a single gamete, 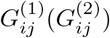, 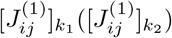 and 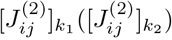 are mixed via row vectors with equal probabilities from *k*_1_(*k*_2_). In RO, which was investigated as a comparison, there are two parents (parent−1 and parent−2). The *i*-th row of both the 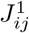 and 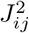 matrices in the next generation are generated from the *i*-th row of those from one of the parents. In contrast to MO or MR, this protocol does not make gametes and genomes in the *i*-th row of the diploid are not mixed. This last case is biologically unrealistic and was added purely to examine the importance of meiosis.

In all four cases, mutation is added after the offspring are produced. Mutations are implemented by renewing one connection of network 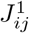 or 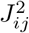 by +1, −1, or 0 according to the probability *μ* per edge(mutation rate).

The evolutionary simulation was conducted using a mutation rate(*μ*) of [10^−5^,10^0^], with the standard deviation of noise(*σ*) being [10^−5^,10^0^]. The number of genes (*N*) was set at 20, the ratio of non-zero elements in *J_ij_* was 0.5, the selection pressure *S* was set at 1, and the number of target genes (*M*) was 5. The relaxation time to test the fitness (T) was 30.

### Group Mendelian ratio

The GMR was computed by adapting the following procedure:

#### 1. Making Homozygous

After evolution, the individuals are usually heterozygotes (e.g., *J1J2*) because it is rare that two genomes are completely identical. Homozygotes are made using the complete copies (e.g., *J1J1* and *J2J2*).

#### 2. Rename haplotypes

The phenotypes *x_i_*(*T*) in one gene *i* of the homozygote are computed to check whether *x_i_* ≈ 1 or *x_i_* ≈ −1. Accordingly, the phenotypes are divided into + or − (e.g., *J1J1* expresses the + phenotype). Note that these signs do not indicate dominance or recessiveness. Haplotypes are renamed according to which phenotype is expressed in the homozygote, + or − (e.g., *J1* is renamed by *+1*).

**Fig 7.**
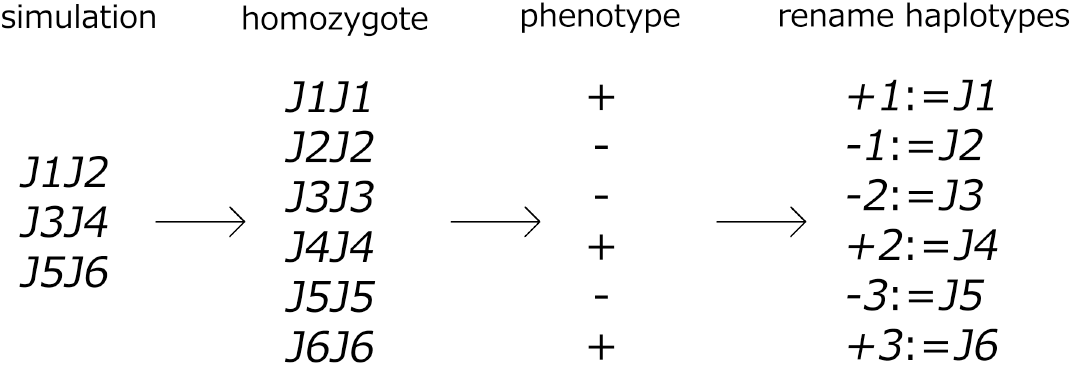
Renaming of haplotypes.

#### 3. Forming a haplotype set to test the “homo” and hetero cases

Haplotype sets are defined as a set of genotypes that express the same phenotype (e.g., *+1*, *+2*, …). To test Mendel’s law, we made two haplotype sets, + and −. From these, we made sufficient “homo” and hetero samples to test whether Mendelian dominance was achieved.

To test the homo case, *N*_samples_ of “homo” diploids were generated from one haplotype set by choosing two haplotypes at random. Note that the “homo” diploid here does not necessarily contain the same *J_ij_*. Then, the phenotypes of the “homo” diploids were computed. The number of samples whose phenotypes were the same as the phenotype of the original haplotype set was counted. This number was defined as 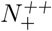 and 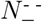 for the phenotypes + and −, respectively.

To test the hetero case, hetero diploids are made by two haplotypes chosen from different haplotype sets. We chose 2*N*_samples_ of the hetero diploid to perform a rigorous comparison with classic Mendelian dominance. By computing the phenotype of each hetero diploid, we obtained the number of dominant phenotypes, which was defined as 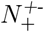 when + is dominant. Here, the samples producing the dominant phenotype were defined as the phenotype that appears most frequently in the hetero diploid result.

#### 4. Calculating the group Mendelian ratio

Following the Mendelian dominance, the GMR was computed by

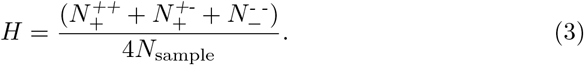

When the GMR is 1, complete GMD is achieved. Conversely, when the GMR = 0.5, the phenotype is determined at random without any dominance.

**Fig 8.**
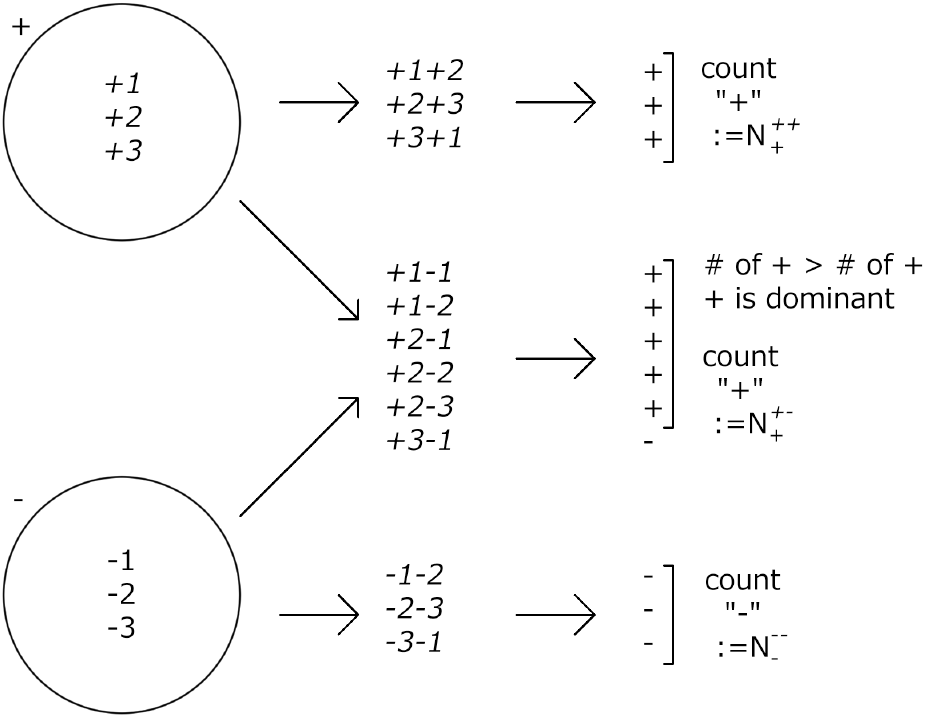
Testing the homo and hetero cases.

### Robustness against initial noise

We computed the average fitness with 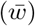 and without 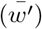 initial noise. Then, the robustness to the initial noise was calculated as the ratio 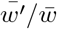. Here, this initial noise was added to swap the sign of *x_i_*(0), i.e., *x_i_*(0) = +1 ↔ −1.

### Distance between diploids

To measure genetic variance, the distance between two genomes was calculated. The distance 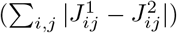 was calculated for two genomes, 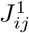 and 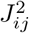, in the diploid. The reported distance was calculated the average distance over all individuals.

**Fig. S1 Diploid distance as a function of mutation rate (including the recombination only case).**

## Acknowledgments

The authors would like to thank Tetsuhiro Hatakeyama and Nobuto Takeuchi for stimulating discussions. This research was partially supported by a Grant-in-Aid for Scientific Research (A) (no. 20H00123) and a Grant-in-Aid for Scientific Research on Innovative Areas (no. 17H06386) from the Ministry of Education, Culture, Sports, Science and Technology (MEXT) of Japan.

